# Benchmarking cell-type deconvolution in cross-platform transcriptomic data

**DOI:** 10.1101/2025.11.24.690141

**Authors:** Aakanksha Singh, Pinar Cakmak, Jennifer H. Lun, Jadranka Macas, Karl H. Plate, Yvonne Reiss, Jonathan Schupp, Katharina Imkeller

## Abstract

**Background:** Transcriptomic data from diverse measurement technologies are widely used to study tissue heterogeneity. Cell-type deconvolution, which resolves mixed transcriptomic signals into cellular components, is a key analytical approach. However, achieving accurate deconvolution across platforms remains challenging due to platform-specific experimental and technological biases.

**Results:** We systematically benchmarked deconvolution performance using real-world cross-platform datasets and simulated data modeling distinct technological features. Our analyses provide practical guidelines for experimental design and promote more robust and comparable cross-platform transcriptomic analyses.

**Conclusion:** Log-normal regression methods such as SpatialDecon demonstrated the most reliable and consistent performance across both simulated and experimental settings, establishing a robust framework for accurate cross-platform transcriptomic deconvolution. Moreover, our results highlight potential caveats in deconvolution predictions arising from specific experimental conditions, providing guidance for more informed experimental design and interpretation

**Graphical Abstract:** 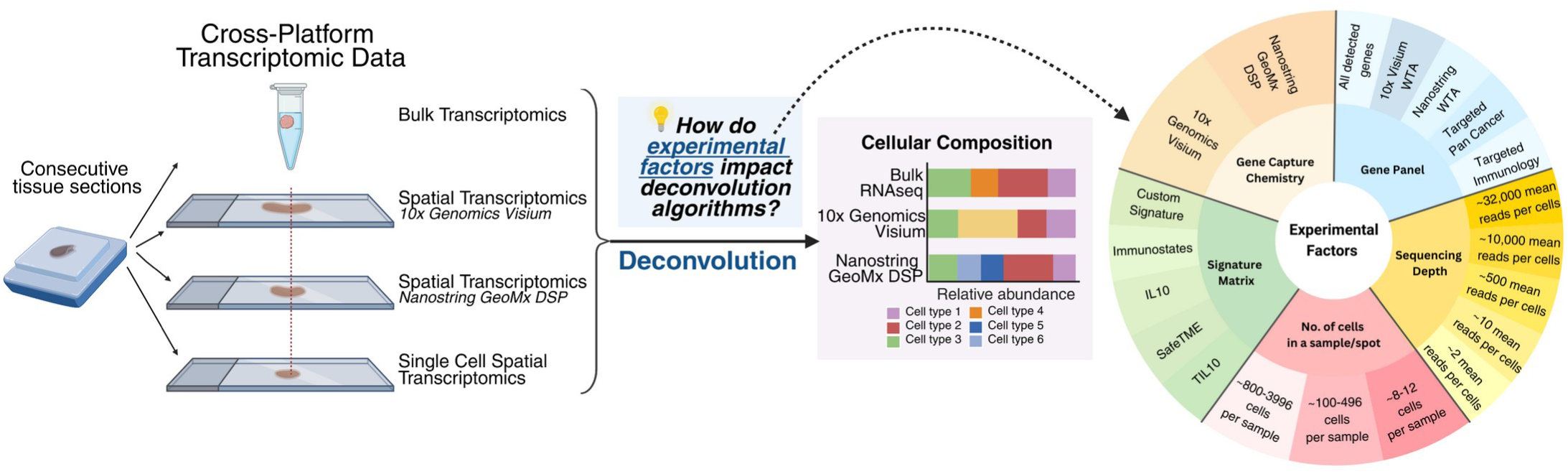

## INTRODUCTION

With rapid technological advances, researchers can now quantify gene expression in biological samples using a wide range of transcriptomic profiling platforms. These include bulk transcriptomics from homogenized tissue, single-cell transcriptomics (e.g., 10x Genomics Chromium, BD Rhapsody), spatial transcriptomics (e.g., spot-based 10x Genomics Visium, region-based Nanostring GeoMx DSP), and spatial single-cell transcriptomics (e.g., 10x Genomics Xenium, NanoString CosMx). These technologies differ in resolution, capture chemistry, tissue processing requirements, number of target genes and number of cells profiled. Despite these differences, all of them generate a count matrix that quantifies the gene expression across samples, spots or individual cells.

To balance cost with measurement detail and to address diverse biological questions, studies increasingly integrate transcriptomic profiles across multiple platforms^1–4^. For instance, a single tissue specimen may be analysed using different technologies, for example bulk transcriptomic and region-based spatial transcriptomics, to gain complementary information. In other cases, publicly available datasets produced using different transcriptomic profiling methods are integrated to investigate specific tissue type or disease condition. Longitudinal samples may be profiled using the same technology but with different chemistries or updated kit versions. Although such scenarios are increasingly frequent, there is currently little consensus on how to consistently and reliably compare transcriptional profiles and downstream analyses across different transcriptomic platforms.

Transcriptional cell-type deconvolution is a common downstream analysis method for datasets without single-cell resolution. It computationally separates a bulk gene expression matrix into its constituent cell types. This approach is particularly valuable for estimating immune cell composition within complex tissues, with applications in oncology and immunology^5–7^. Table 1 lists commonly used computational tools for estimating cell type abundances across different transcriptomic data types.

**Table 1:** Overview of transcriptional deconvolution methods.

| Tool | Designed for | Mathematical model | Signature matrix customizable? |
| --- | --- | --- | --- |
| MCPcounter<sup>2</sup><sub>2</sub> | Signature quantification, bulk | log-sum of marker gene expression | no |
| CIBERSORT<sup>39</sup> | Deconvolution, bulk | support vector regression | yes |
| EPIC<sup>21</sup> | Deconvolution, bulk | constrained least squares regression | yes |
| quanTIseq<sup>20</sup> | Deconvolution, bulk | constrained least squares regression | yes |
| SPOTlight<sup>32</sup> | Deconvolution, spot-based spatial | seeded non-negative matrix factorization (NMF) | requires a mandatory single-cell dataset to build reference |
| Celloscope<sup>54</sup> | Deconvolution,<br>spot-based spatial | Bayesian probabilistic graphical<br>model | yes |
| CARD<sup>55</sup> | Deconvolution,<br>spot-based spatial | conditional autoregressive<br>model-based deconvolution | requires a<br>mandatory single-cell dataset to<br>build reference |
| cell2location<sup>5</sup><br>6 | Deconvolution,<br>spot-based spatial | Bayesian mixture model | requires a<br>mandatory single-cell dataset to<br>build reference |
| SpatialDecon<sup>23</sup> | Deconvolution,<br>region-based spatial | log-normal regression | yes |

\* MCPcounter outputs are most suited for comparing a given cell type across samples (Becht et al., 2016), but not for comparing different cell types within a sample. In this study, scores were scaled to the interval [0,1] to generate relative cell-type specific value that preserves across-sample distributions (see methods).

Multiple benchmarking studies have evaluated transcriptional deconvolution tools for both bulk^8–13^ and spatial data^14–16^, focusing primarily on metrics such as accuracy and computational efficiency. However, most benchmarks have not considered how varying experimental conditions and technological differences across transcriptomic platforms influence the performance and reliability of deconvolution algorithms.

In this study, we evaluate whether deconvolution tools produce reliable and reproducible results across different transcriptomic platforms and experimental conditions. To address this, we apply multiple deconvolution methodologies to real-world cross-platform transcriptomic datasets.

Additionally, we generate simulated datasets to disentangle the effects of specific technological factors. We systematically investigate how the following experimental factors, known to differ across transcriptomic platforms, affect deconvolution outcomes: (i) Gene capture chemistry: The underlying technological principle used to measure gene expression. For example, 10x Visium v1 3’ captures transcripts through poly-A tail binding, whereas 10x Visium v2 and the Nanostring GeoMx DSP relies on barcoded probes that hybridize to target mRNA. (ii) Gene panel: The specific set of genes covered by a transcriptomic platform. The gene panel may cover the whole transcriptome (whole transcriptome assay, WTA) or a pre-defined subset of genes (e.g., 10x Genomics Visium targeted Pan Cancer and targeted immunology panels). (iii) Sequencing depth: The number of sequencing reads measured per cell in the region of interest or sample. This measure is particularly relevant for WTAs, since the sequencing depth also determines the number of genes detected. (iv) Number of cells measured: The cellular resolution of the assay. Bulk transcriptomics cover ca. 10^3^-10^6^ cells, whereas region-based spatial methods such as Nanostring GeoMX ROIs cover ca. 10^2^-10^3^ cells^17^ and spot based spatial methods such as 10x Visium (v2, v1 3’) only cover ca. 10 cells per spot^18,19^.

Our results highlight how the different mathematical approaches used in the deconvolution tools are affected by data properties that depend on technological differences between transcriptomic platforms. Moreover, we provide practical guidelines computational method choice dependent on experimental design for robust deconvolution of cross-platform transcriptomic data.

## RESULTS

### Transcriptional deconvolution results vary across transcriptomic platform and deconvolution tool

Many recent biological studies integrate transcriptomic data from multiple technological platforms to address different biological questions, to balance cost against resolution, or to leverage existing public datasets. To illustrate the challenges of cross-platform transcriptomics and their impact on transcriptional deconvolution and biological interpretation, we reanalysed a dataset from one of our previous studies in which consecutive tissue sections were profiled using different omic methods. We include the following datasets focusing on immune aggregates in brain tumor tissue (Fig. 1A): Multiplex Immunofluorescence (mIF)^1^: 10 glioma FFPE blocks were stained for B cells (CD20) and T cells (CD3). A pathologist annotated aggregates of immune cells and counted the number of B and T cells in each immune aggregate to establish ground truth cell counts from proteomics assay. Nanostring GeoMX DSP^1^: Consecutive tissue sections from the FFPE blocks were obtained and 28 of the immune aggregates identified in mIF were further profiled using the Nanostring GeoMX DSP platform (WTA gene panel). Immune aggregates from mIF and Nanostring GeoMX areas can directly be mapped to each other, allowing direct comparison of immune cell frequencies between transcriptomics and proteomics assays. 10x Visium (generated in this study): For a subset of glioma (n = 2) and brain metastasis (n = 2) patients, we performed an additional 10x Genomics Visium v2 FFPE WTA assay on consecutive tissue sections. After sequencing, data corresponding to spots within CD20-enriched regions were extracted (approximately 50–100 spots per patient) following visual inspection of CD20 gene expression using the Loupe Browser. These regions were considered immune aggregates.

**Figure 1:**
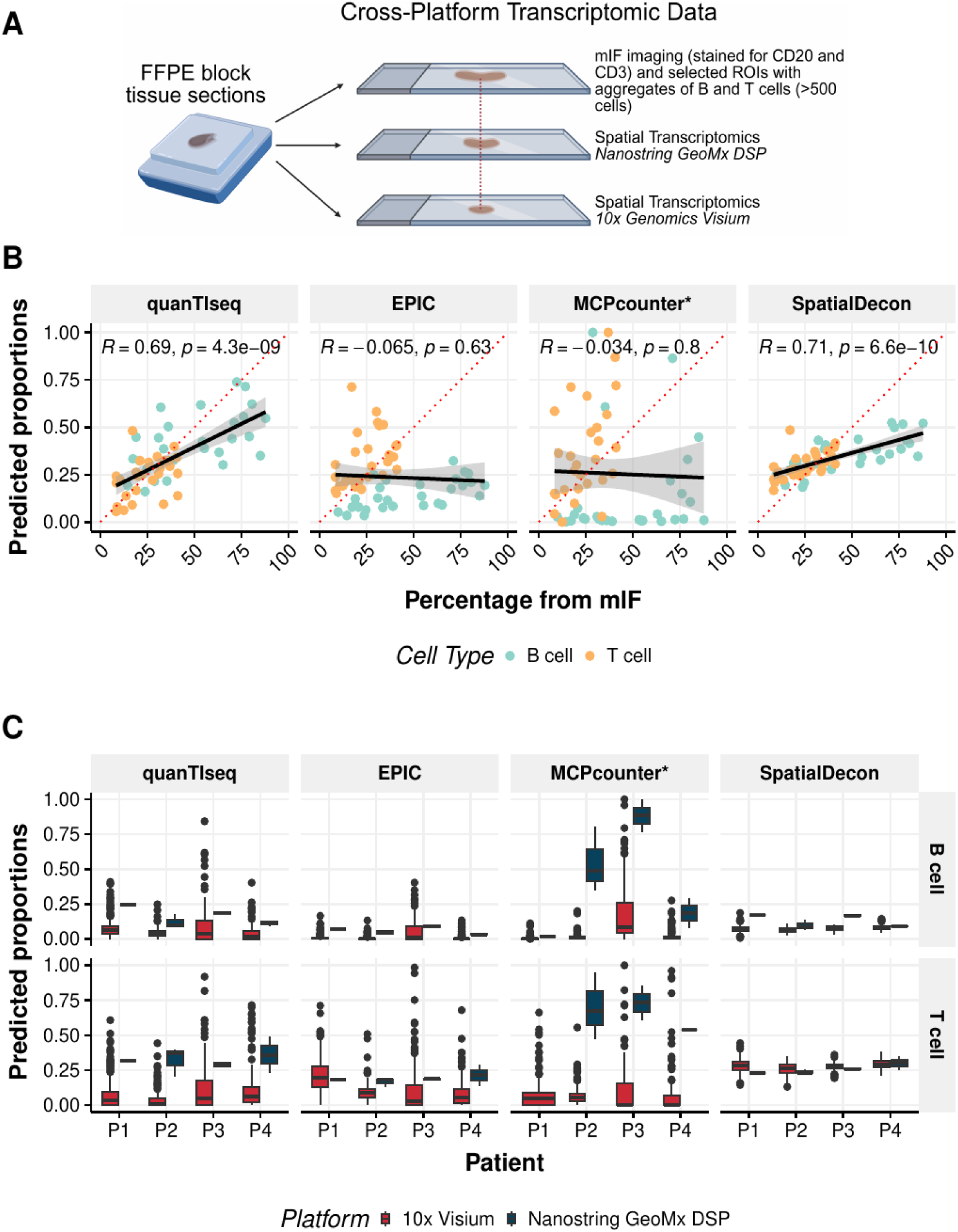
Quantification of B and T cells in cross-platform transcriptomic data. **(A)** Schematic representation of matched regions of an FFPE tissue block profiled by mIF (stained for CD20 and CD3), Nanostring GeoMx DSP, and 10x Genomics Visium v2 FFPE spatial transcriptomics. **(B)** PCC between proportions of B and T cells as seen in mIF images (stained for CD20 and CD3) and the predicted proportions of the cells in the consecutive tissue section that was processed using Nanostring GeoMX DSP. The red dotted line represents the line of identity (x = y). *MCPcounter scores were scaled to [0,1] to create relative cell-type values while preserving their distribution (see methods). **(C)** Predicted proportions of B and T cells from regions with immune cell aggregates from 10x Visium spots and Nanostring GeoMX DSP ROIs from four patients. Boxplots represent median, 25th and 75th percentiles. There are multiple spots (ca. 50-100) in the areas designated as ‘immune cell rich’ in 10x Visium which are compared against 1-4 regions of interest in Nanostring GeoMx DSP per patient.

We first compared frequencies of B and T cells extracted from mIF with those predicted by deconvolution tools applied to the GeoMx DSP data (Fig. 1B). Deconvolution was performed using quanTIseq^20^, EPIC^21^, MCPcounter^22^, and SpatialDecon^23^. These methods represent a subset of deconvolution methods that reflect a broad range of design choices and underlying mathematical models (Table 1). We initially used each method’s default signatures to reflect common user practice and provide a pragmatic baseline for cross-platform users. Across 28 ROIs with matched mIF data the overall correlations were moderate and varied across tools. SpatialDecon consistently achieved the highest concordance (Pearson correlation coefficient (PCC) = 0.71) between predicted and observed B and T cell proportions, followed closely by quanTIseq (PCC = 0.69).

Next, we compared B and T cell proportions predicted from GeoMx ROIs and Visium spots (Fig. 1C). In this analysis, we were not able to directly match the regions of interest accross assays due to considerable histomorphological differences between the consecutive tissue sections. However, we were able to compare the combined regions of immune aggregates originating from the same tissue block. For this, deconvolution was performed on the ROI level for GeoMx DSP data and on spot level for the Visium spots (immune aggregates delienated manually using B and T cell marker genes). Although the cellular composition of immune aggregates is expected to be comparable within a given patient^1^, predicted proportions differed substantially between the two platforms, with the extent of difference in predictions depending on the deconvolution method used (Fig. 1C). Least-square regression methods (EPIC, quanTIseq) systematically estimated higher B and T cell proportions in GeoMx DSP data compared to 10x Visium data. Among the tested tools, SpatialDecon showed smallest differences between mean predicted proportions but also the small variability in predicitons within one platform.

### Gene panel selection impacts deconvolution results in simulated and experimental data

Given the large variability in deconvolution results across computational methods and technological platforms, we next investigated the impact of individual experimental factors that differ between technologies. We began by examining the effect of the gene panel, i.e. the set of genes measured in a transcriptomic assay. Pre-designed targeted gene panels, which capture only a fraction of the whole transcriptome, are often used to balance cost and technological constraints while capturing the most informative genes. Since gene panels vary widely in size and gene selection, they are expected to influence deconvolution outcomes, with the magnitude of this effect likely depending on the method used.

To comprehensively evaluate the choice of the gene panel on the deconvolution results, a systematic data simulation approach was implemented (Fig. 2A). For this, we used single-cell RNA-seq datasets of PBMCs (Dataset A, B, C and D listed in Table S1) and healthy bone marrow (Dataset E, F and G listed in Table S1). We annotated various immune cell types, including B cells, T cells, monocytes, and NK cells, dendritic cells, basophils, neutrophils and progenitors using the Monaco Immune Data^24,25^ as a reference and the SingleR package^26^ for automated cell-type classification (Table S2). We then created pseudo-bulk samples by aggregating counts from B cells, T cells, monocytes, and NK cells that were randomly sampled from the annotated cell pool of each cell type in a dataset to generate a pseudo-bulk-level count matrix.

**Figure 2:**
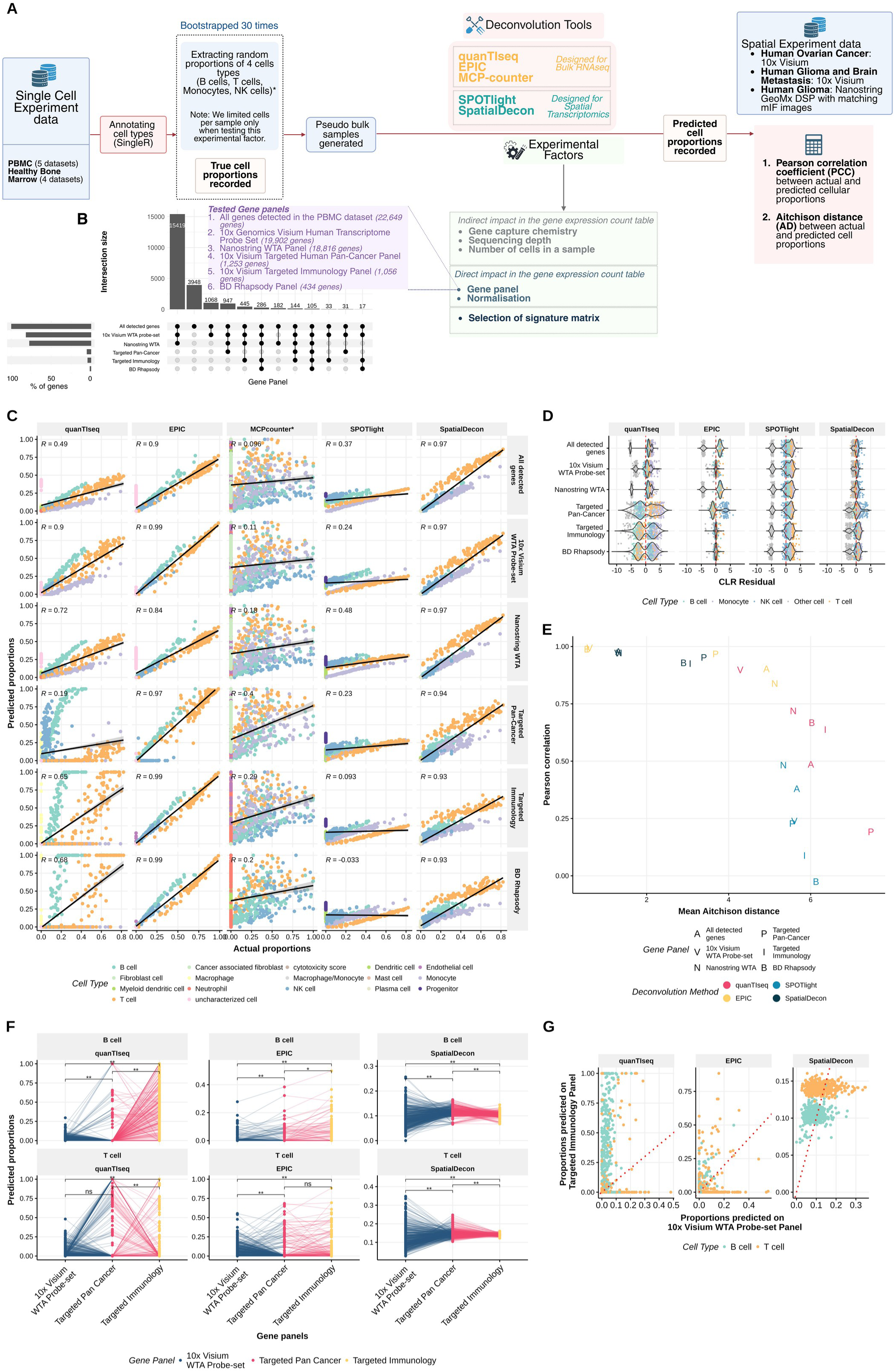
Impact of gene panel on deconvolution performance. **(A)** Schematic representation of the approach used to simulate pseudo-bulk samples from publicly available PBMC and bone marrow single cell datasets. When testing a particular experimental factor, all other factors were kept constant using their default settings. In this analysis, we tested five commonly used gene panels across five deconvolution tools. **(B)** Upset plot showing gene distribution across five gene panels. Three panels (all genes, 10x Genomics, and Nanostring WTA Panel), are large, each containing over 15,000 genes. In contrast, BD Rhapsody Panel and the 10x Genomics Xenium Human Immuno-Oncology panel are much smaller, each representing less than 1% of the total number of genes. **(C)** Correlation plots comparing actual and predicted cell type proportions in PBMC pseudo-bulk samples. Points are colored according to cell type. Each plot facet displays R, which is the PCC. **(D)** Distribution of CLR residuals across deconvolution methods and gene panels. The color of the dots denotes the cell type. **(E)** Performance relationship between PCC and mean AD across gene panels on the deconvolution tools in the PBMC pseudo-bulk samples. **(F)** Predicted proportions of B and T cells across 500 randomly selected spatial spots from the same tissue section in the Human Ovarian Cancer dataset (10x Visium), evaluated across three gene panels. Proportions range from 0 to 1. Pairwise Wilcoxon tests were performed between panels, with Bonferroni correction applied. Asterisks indicate significance: p = 3.30 × 10⁻² for the comparison between the Targeted Immunology and Targeted Pan-Cancer panels using EPIC. All other comparisons marked with “**” have p < 1.5 × 10⁻¹⁴. **(G)** Predicted B and T cell proportions for 500 randomly selected spatial spots in the human ovarian cancer dataset, comparing the Targeted Immunology panel to the 10x Visium WTA panel. Each dot represents the same spot analysed with both panels. Colored dots represent B cells and T cells. R indicates the Pearson correlation coefficient (PCC) between panels for each cell type. The red dotted line represents the line of identity (x = y).

The genes in this matrix were filtered to match four transcriptomic panels: BD Rhapsody Immune Response (in-house, 434 genes)^27^, targeted immunology^28^ (1,056 genes), targeted pan-cancer^29^ (1,253 genes), Nanostring GeoMx WTA^30^ (18,816) and 10x Visium Human Transcriptome^31^ (19,902 genes). In addition, we included a panel containing all genes detected in the dataset with existing HUGO gene symbol that were detected in the dataset (22,649 genes in PBMC and 33,514 genes in Healthy Bone Marrow; see Fig. 2B). A full list of genes included in each gene panel is provided in Table S3. We applied four deconvolution tools (quanTIseq, EPIC, MCPcounter, SpatialDecon), using their default signature matrices, as well as SPOTlight^32^ with Dataset D (Table S1) as signature reference. Predicted cell proportions were then compared to the ground truth across all simulated datasets.

We first evaluated deconvolution performance using PCC between the predicted and actual cell proportions (Fig. 2C). However, compositional data such as cell-type proportions are susceptible to spurious correlations^33^. To address this, we used the residuals from the centered log-ratio (CLR) transformation to quantify the differences between actual and predicted cell-type compositions in a pseudo-bulk sample^34^ (Fig. 2D). Based on these CLR residuals, we calculated the Aitchison distance (AD) for each sample, which measures the dissimilarity between the compositional vectors of actual and predicted cell type proportion^35^ (Fig. 2E) (see Materials and Methods for details). Across all gene panels and for both PBMC and bone marrow datasets, EPIC followed by SpatialDecon performed most consistently, showing the highest PCC, the lowest absolute CLR residuals and lowest mean AD (Fig. 2C to E, Fig S1). Of note, the default implementation of EPIC does not support monocyte prediction, so monocyte were excluded from all EPIC-related analyses including the generation of simulated data.

The two constrained least square regression methods, quanTIseq and EPIC performed well on large panels. Interestingly, both tools performed best with the 10x Visium WTA panel, and less well with the Nanostring WTA panel and the complete WTA panel (Fig. 2C, D and Fig. S1A, B). This decreased in performance was mainly due to the prediction of a high proportion of “uncharacterized cells”, which are a result of gene expression that cannot be explained by a function of the signature matrix. This highlights that the prediction of uncharacterized cells by constrained least square regression methods is sensitive to gene panel selection.

The performance of quanTIseq decreased for smaller gene panels (Fig. 2C-E). EPIC failed to predict NK cells for the Targeted Pan-cancer gene panel (Fig. 2C-D) These effects likely stems from the different signature matrices used in both tools. In particular, quanTIseq had minimal overlap with some panels, e.g. only 23 genes are common between the targeted pan-cancer and the default quanTIseq signature matrix.

SPOTlight showed poor performance across all gene panels in PBMCs, with PCC values less than 0.5 (Fig. 1D), and negative values of PCC in bone marrow samples (Fig. S1A). The tool tend to overestimate the proportions of progenitors and dendritic cells in the larger gene panels, at the expense of monocytes. Simiarly in the smaller gene panels, it also misattributed proportions of monocytes and T cells (Fig. 2D and Fig. S1).

The comparisons of MCPcounter results with all other tools are more challenging, as it is based on gene signature quantification. To enable comparison, we scaled both the actual proportions and MCPcounter scores to the interval [0,1] for each cell type, thereby preserving the relative distribution across samples (for details see methods). As a result, the normalized scores did not preserve compositionality within a sample and we could therefore not calculate CLR residuals and AD for MCP-counter. MCPcounter failed to detect of small differences in cell type abundance across samples (Fig. 1C).

We next aimed to validate the differences in deconvolution performance across tools and gene panels using real-world cross-platform transcriptomic data. For this, we analysed publicly available 10x Visium spatial transcriptomic data from the same section of human ovarian cancer tissue, for which three different sequencing libraries covering different gene panels were prepared: whole transcriptome (WTA), targeted immunology (TI; 1,056 genes), and targeted pan-cancer panel (TPC; 1,253 genes) (see Table S1). We applied three deconvolution methods (quanTIseq, EPIC, and SpatialDecon) and focussed on the estimated B and T cell proportions across 500 spatial spots (Fig. 2F). Predicted cell proportions for quanTIseq and EPIC varied notably for the same spot, particularly as gene panel size decreased (Fig. 2F,G). For example, quanTIseq predicted significantly higher B and T cell proportions with the ‘Targeted Immunology’ panel than the other two panels (Fig. 2F, G), which is also reflective of the observations made with simulated data. Despite EPIC’s good performance on simulated data, the tool did not achieve notable correlation between predictions on WTA and Targeted immunology panel in the experimental data (Fig. 2G). SpatialDecon showed the greatest variation in B and T cell predictions across spots with the ‘WTA’ panel, but this variation was nearly absent with the ‘Targeted Immunology’ panel (Fig 2F, G). This decreased variance in predicted values with decreasing gene panel size was also observed in the deconvolution of simulated data. With decreasing gene panel size, least square regression techniques, in particular quanTIseq with default signature matrix, tend to overestimate variability between samples, whereas log-normal regression techniques such as SpatialDecon underestimate this variability.

### Commonly used sequencing depths have minimal effect on deconvolution tools

Another important experimental factor that affects transcriptomic profiles across technological platforms and experimental designs is sequencing depth. To evaluate how the average number of reads per cell affects deconvolution accuracy, we randomly downsampled reads from FASTQ files of two single-cell datasets. PBMC (Dataset H; Table S1, Fig 3) and bone marrow (Dataset I; Table S1, Fig S2) datasets were downsampled to 60%, 20%, 1%, 0.02% and 0.002% of the original reads (Fig. 3A and Fig. S2). Cell Ranger^37^ 9.0.0 was reran for each subsampled dataset to generate filtered feature-barcode matrices. The number of cells for each subsampled single cell dataset was relatively stable down to 1% of original reads. To ensure accurate cell-type annotation, we first annotated cells using the full dataset (100% reads) and transferred these labels to the downsampled datasets, as annotation becomes unreliable at very low read depths. We then generated pseudo-bulk samples by aggregating cells per sample, using the workflow described in Fig. 2A.

**Figure 3:**
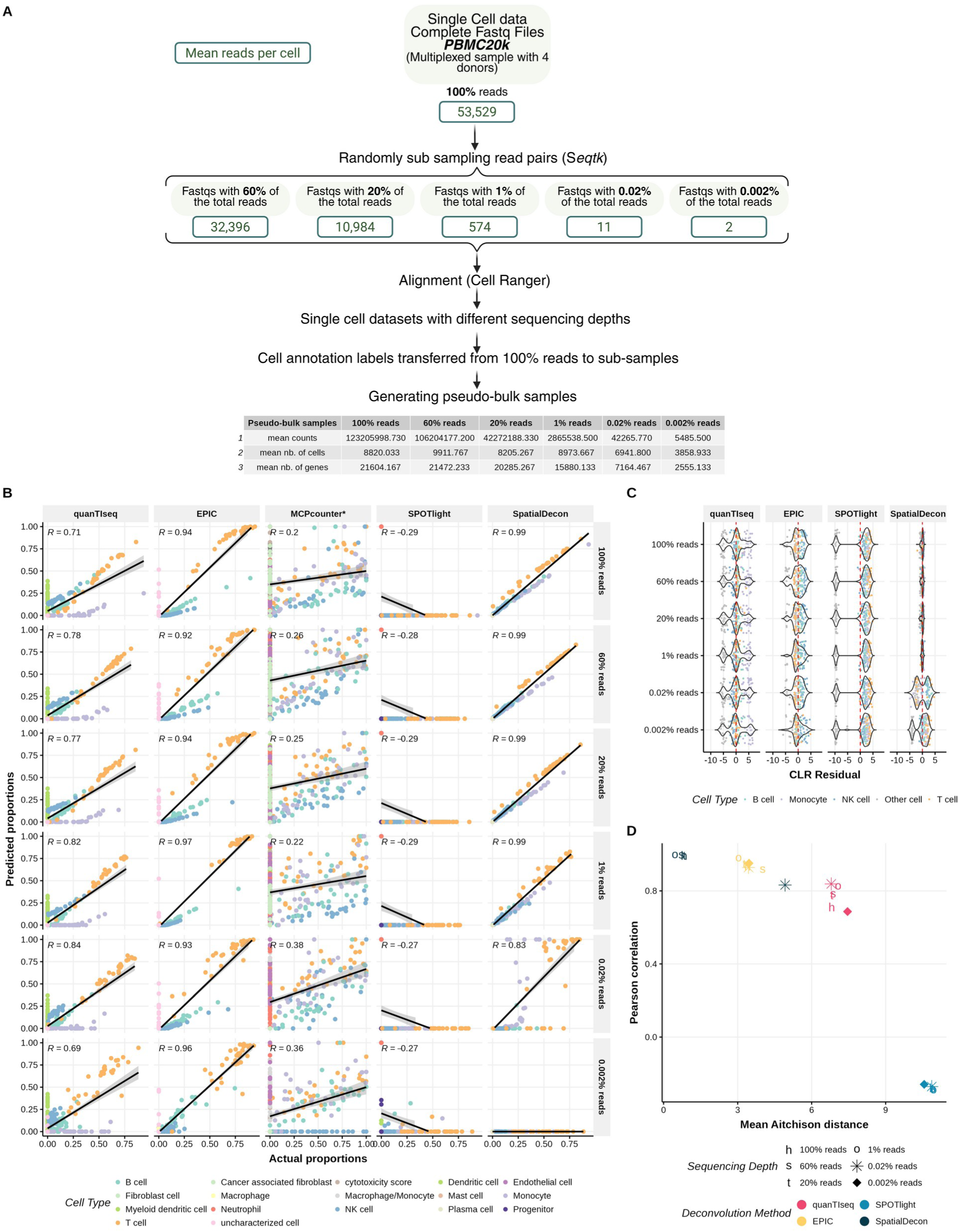
Impact of sequencing depth the deconvolution performance in PBMC pseudo-bulk samples. **(A)** Reads were randomly downsampled from FASTQ files to obtain 100%, 60%, 20%, 1%, and 0.02% and 0.002% of the original read depth. For each subsample, CellRanger was re-run and cell annotations were transferred from the full dataset (100% reads). Pseudo-bulk samples were generated using the workflow described in Fig. 1B. **(B)** Correlation between actual and predicted cell type proportions in PBMC pseudo-bulk samples across sequencing depths. The colors of the dots represent the different cell types. PCC is shown in each plot. **(C)** Distribution of the CLR residuals for each deconvolution method at different sequencing depths. The color of the dots denotes the cell type. **(D)** PCC versus mean AD for the performance across sequencing depths and deconvolution methods.

Contrary to expectations, the PCC did not decrease with reduced sequencing depth and the distribution of CLR residuals and AD remained stable across sequencing depths down to 1% of total reads (Fig. 3B to D, Fig. S2). While SpatialDecon performance declined below 1% sequencing depth (∼574 reads/cell for PBMC, Fig 2A; ∼399 reads/cell for bone marrow, Fig S2A), quanTIseq and EPIC showed minimal deterioration even at 0.02% sequencing depth of (∼11 reads/cell for PMBC, Fig 3A; ∼9 reads/cell for bone marrow, Fig S2A). These findings suggest that sequencing depth can be reduced substantially, to a few hundreds of reads per cell, without significantly affecting deconvolution accurancy, offering a potential cost-saving strategy. Constrained least square regression methods seem to deliver more reliable results in samples with low sequencing depth (Fig. 3B to D).

### Impact of the number of cells per sample or spot on deconvolution performance

Transcriptomic platforms capture gene expression at varying cellular resolutions, ranging from spot-based spatial transcriptomics (5–18 cells per spot), to region-based spatial transcriptomics (hundreds to thousands of cells per region), up to bulk transcriptomics (thousands to millions of cells per sample). Accordingly, deconvolution tools have been developed with different assumptions regarding the number of cells per measurement. For example, SPOTlight was developed for small spatial transcriptomic spots and leverages information across multiple spots, whereas SpatialDecon was designed for GeoMX DSP regions that typically contain hundreds of cells. In contrast, bulk-oriented tools such as quanTIseq, EPIC, and MCP-counter were developed for samples containing thousands to millions of cells.

In our previous simulations, we used an average of ∼2,000 cells per sample in the PBMC dataset and ∼600 cells per sample in healthy bone marrow datasets, mimicking bulk-like conditions. To systematically assess how input cell number affects deconvolution performance, we simulated pseudo-bulk samples composed of four (except EPIC, where we simulated only 3 cell-types) cell types at varying total cell counts: 600-3996 cells per sample (200–999 of each type), 75-496 cells per sample (25–124 of each type), and as few as 6-12 cells per sample (2–3 of each type) (Fig. 4A). Because of these constraints, the true proportion of any individual cell type ranged from 0.06 to 0.7 in these simulations.

**Figure 4:**
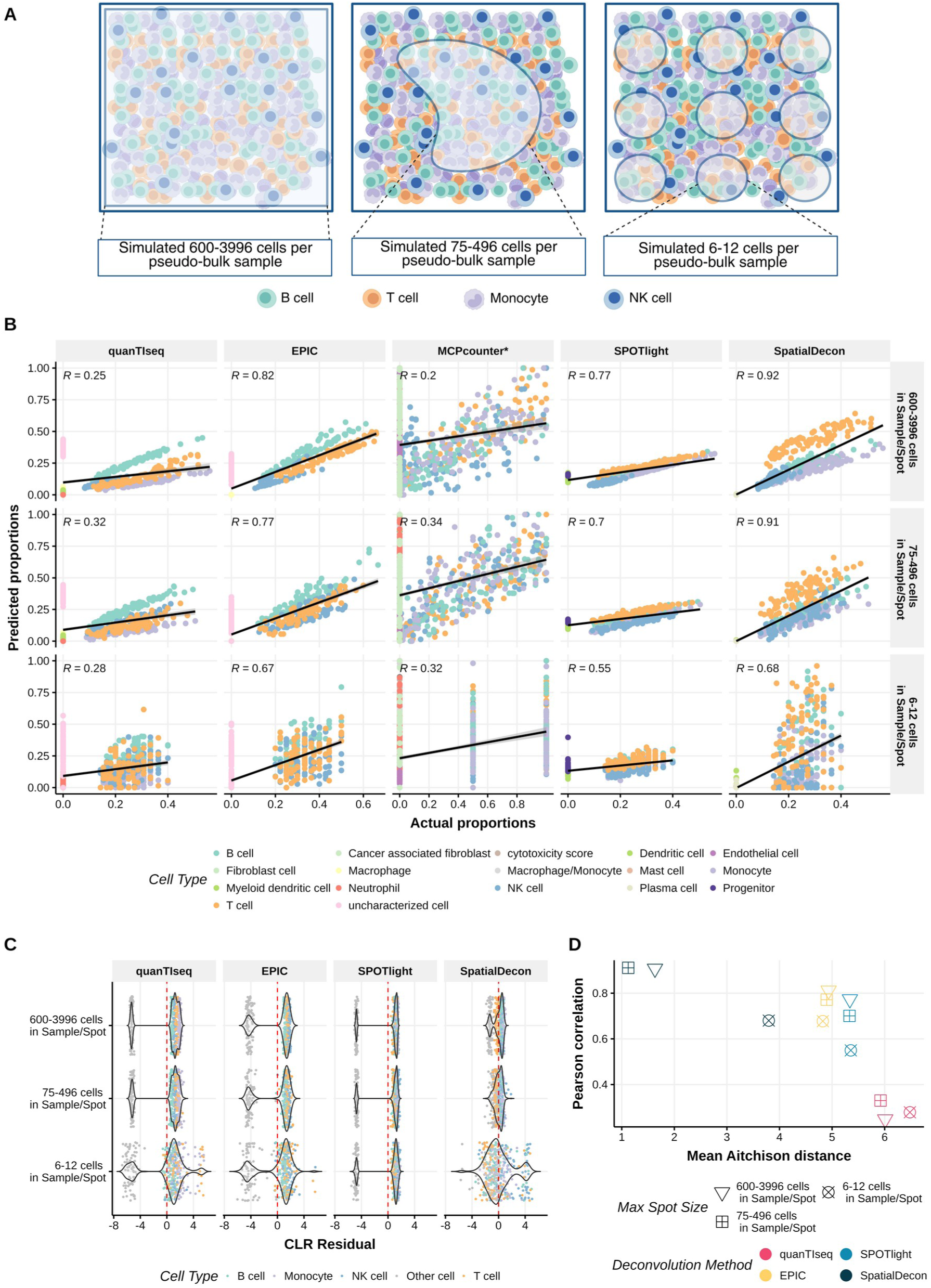
Impact of the total cell number on deconvolution performance. **(A)** Schematic representations of simulating pseudo-bulk samples or spots of varying sizes. Maximum cell numbers were restricted to 4000, 500, and 12 cells, simulate pseudo-bulk samples resembling bulk RNA-seq, Nanostring GeoMx DSP region of interest (ROIs) and 10x Visium spatial transcriptomic spots, respectively. **(B)** Correlations between actual and predicted cell type proportions in PBMC pseudobulk samples. The colors of the dots represent the different cell types. The PCC, R, is shown in each facet. **(C)** Distribution of the CLR residuals for the four cell types included in the PBMC pseudo-bulk samples. Each dot represents a cell type and is colour-coded. **(D)** PCC versus mean AD, summarising the performance of each deconvolution tool across sample sizes.

Highest differences between actual and predicted proportion and highest prediction variability was observed for log-normal and least-square regression methods in the smallest cell number setting (Fig. 4B-D). Across all settings, SpatialDecon demonstrated the highest accuracy with strong correlation and low AD, followed by EPIC (Fig. 4B-D). SPOTlight performance was least affected by reducing the number of cells in a sample (Fig. 4B-D), which reflects the superiority of NMF-based models to estimate cell frequencies in very small samples.

### Gene count normalisation does not affect previous deconvolution results

Normalisation is a standard preprocessing step to account for differences in sequencing depth or sample size. In our previous analyses, we applied TPM normalisation (similar to library-size normalization) prior to all deconvolution approaches, independent of a tool’s documentation that recommended an alternative strategy. To ensure this choice did not bias our results, we tested each deconvolution method in combination with three commonly used normalisation strategies: TPM normalisation, Q3 normalisation, and quantile normalisation (Bolstad et al., 2003) (Fig. S3A). Across both the PBMC and bone marrow datasets, SpatialDecon results produced consistently high PCC results across all normalisation approaches (Fig. S3B, C, E, F). In contrast, quanTIseq and EPIC showed reduced performance with Q3 normalisation, while TPM, raw, and quantile normalisation yield comparable outcomes. Overall, normalisation only had marginal influence on performance within each tool, and importantly, the use of TPM normalisation in our prior analyses did not negatively affect predictions (Fig. S3D, G).

### Choice of signature matrix should depend on gene panel and the cells of interest

As a final step in our study, we investigated the impact of the signature matrix, i.e. the cell type–specific gene expression reference used for deconvolution, on deconvolution performance. Signature matrices can be either be pre-defined or custom-built from tissue-specific single-cell data. First-generation deconvolution tools rely on pre-defined references, whereas second generation tools, such as SpatialDecon, allow the incorporation of custom single-cell references^13^. However, in many cases, single-cell datasets from the same tissue may be unavailable.

To assess how well publicly available signature matrices can deconvolute target cell types, we used SpatialDecon, given its robust performance across conditions and support for both built-in and custom matrices. We evaluated four widely used immune cell signature matrices alongside a custom single-cell–derived reference (Dataset D for PBMC and Dataset F for health bone marrow pseudo-bulk samples). The tested matrices included TIL10^20^, ImmunoStates^38^, LM22^39,40^, and safeTME^23^ (Fig. 5A). These signature matrices vary widely in size and gene content, e.g. TIL10 comprises only 170 genes and overlaps with just 9 genes with the others (Fig. 5A).

**Figure 5:**
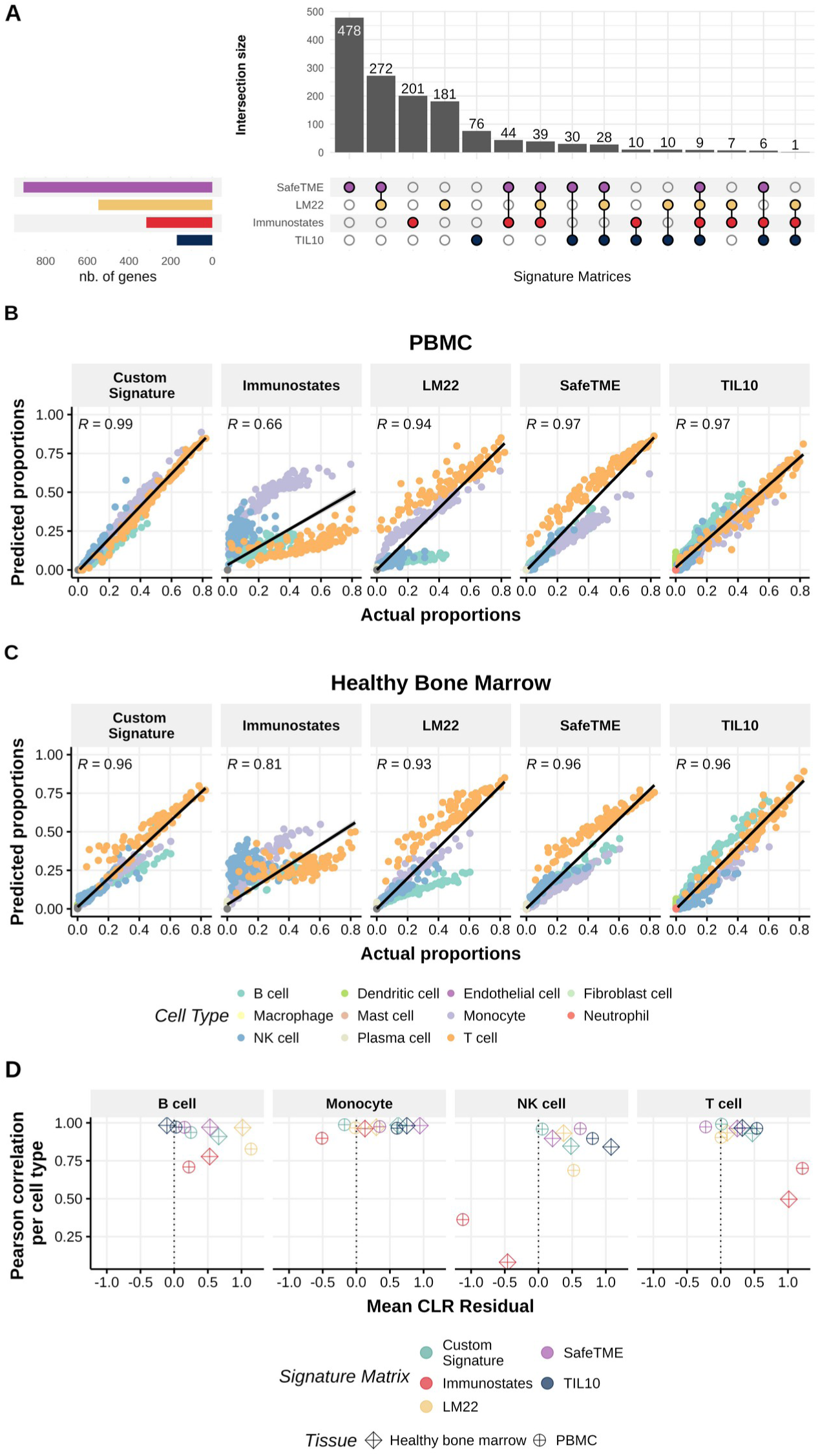
Impact of choice of signature matrix on predicted proportions of cells of interest. **(A)** Upset plot showing gene overlap among four commonly used immune cell signature matrices: TIL10 (default signature matrix used in quanTIseq), ImmunoStates, LM22 (default signature matrix used with the tool CIBERSORT^39^) and SafeTME (default signature used in SpatialDecon). **(B, C)** Correlations between actual and predicted proportions of cells in the (B) PBMC and (C) bone marrow pseudo-bulk samples. The colors of the dots represent the different cell types. **(D)** Performance evaluation of each signature matrix based on PCC versus mean CLR residuals across four cell types in PBMC and bone marrow pseudo-bulk samples.

As expected, the single-cell derived custom signature matrix showed the highest PCC between predicted and actual proportions (Fig. 5B, 5C). However, TIL10 and safeTME also achieved high PCC values in both PBMC and healthy bone marrow datasets, despite their limited marker genes. This suggests that smaller, well curated gene signature may improve results, provided they sufficiently overlap with the gene panel (as seen in quanTIseq performance with smaller gene panel, Fig. 2).

The accuracy of predicted proportions also varied across cell types (Fig. 5D). Notably, TIL10 outperformed the custom signature for B cell abundance predictions, while LM22 performed best for monocytes in healthy bone marrow. ImmunoStates performed better for monocytes and B cells than for NK cells or T cells. These differences emphasize that no single signature matrix is universally optimal; rather, matrix selection should be guided by both the cell types of interest and the biological context of the study. For studies focusing on adaptive immune responses, signature matrices with well-performing B and T cell signatures are suitable (e.g. TIL10 or custom signature matrices).

## DISCUSSION

The rapid development of new transcriptomic platforms and the increasing need to integrate data across technologies underscores the importance of robust deconvolution methods that enable reliable cross-platform comparisons. In this study, we systematically assessed the performance of several deconvolution tools under a range of technological and experimental constraints, using both simulated pseudo-bulk datasets and real-world data from our experiments and publicly available repositories.

Across all analyses on simulated and experimental data, SpatialDecon, which applies log-normal regression, consistently achieved the most accurate and reproducible results. Its advantage likely stems from its ability to model the inherent skewness and heteroscedasticity of gene expression data. The method proved particularly suitable to compare deconvolution results across gene panels and sample sizes, down to cell number of ∼100 cells per sample. Least-squares regression methods such as EPIC and quanTIseq also performed well but were more likely to predict uncharacterized cells. These methods showed robust performance also with low sequencing depth.

EPIC showed considerable better performance on simulated than on experimental data. Of note, EPIC returns two proportion values: mRNAProportions and cellFractions. The documentation recommends using mRNAProportions for benchmarking the tool with in-silico generated samples and cellFractions for true bulk samples^21,36^, whereas other tools do not offer such options. Therefore, we used mRNAProportions for our analyses. This could potentially explain EPIC’s strong performance on simulated data, which is not observed when comparing deconvolution proportions derived from cellFractions to the ground truth cell counts obtained from mIF (Fig. 1B).

To assess cross-platform performance, we compared consecutive tissue sections analyzed with 10x Visium v2 and NanoString GeoMx DSP, using mIF-derived cell frequencies as benchmarks. Among all tools, SpatialDecon and quanTIseq demonstrated the strongest concordance with experimental data. SpatialDecon also maintained the highest consistency across the two platforms, underscoring its suitability for transcriptional deconvolution in cross-technology studies. Differences in mRNA capture mechanisms likely contribute to observed performance variability: poly(A)-based methods perform best with intact RNA, while probe-based or rRNA depletion–based approaches are preferred for FFPE tissues. These technologies further differ in detection mechanisms (light activation for GeoMx DSP vs. barcoded spatial spots for 10x Visium), which may introduce systematic biases. As the sections were not spatially registered, heterogeneity in predictions between tissue sections may also contribute to inter-platform discrepancies, which could partially explain the differences between the assays in Figure 1C.

Using our pseudo-bulk simulation strategy, we further investigated how sequencing depth and spot size influence deconvolution accuracy. Our data simulation strategy, which includes simulation of sequencing depth and spot size expanded upon existing pseudo-bulk simulation methodologies proven reliable for benchmarking deconvolution tools^9,13,43^. The simulation of sequencing depth indicated that reliable deconvolution using log-normal or least-square regression is achievable with approximately ∼300 reads per cell. While this finding covers deconvolution of broad cell types, higher coverage may be necessary for resolving fine cell type definitions. In our study, larger spot sizes resulted in stable estimates across multiple tools, with best results achieved for SpatialDecon and EPIC. The deconvolution results became less accurate for smaller spots, and SpatialDecon performance was most affected by the decrease in spot-size. Transcriptional deconvolution of small cell numbers proved unreliable, which may partially explain the differences observed in Figures 2F and 2G. It is important to note that our sample-size simulations did not include absent or rare cell types; zero-heavy compositions can affect CLR/AD and regression stability, particularly at 6–12 cells per spot.

We further investigated how the breadth of genes coverage affects deconvolution. As expected, performance decreased sharply when the overlap between the gene panel and the signature matrix was small. Our findings from the gene panel comparisons have broader implications for experimental design and gene panel selection. Notably, the widely used TIL10 signature matrix (170 genes) performed exceptionally well on whole-transcriptome data but exhibited limited overlap (<60 genes) with commonly used targeted immunology or pan-cancer panels. This discrepancy arises because many of the most informative genes in TIL10 are not traditional canonical marker genes. These results suggest that including such non-traditional yet informative genes in targeted panels could significantly enhance cell-type resolution, particularly in immune-rich tissues. Designing probes for such genes may therefore be a valuable strategy for future spatial and single-cell transcriptomic studies. Although our simulations and validations across PBMC and bone marrow datasets show consistent patterns, they may not have fully captured the tumor-specific microenvironmental complexity of FFPE or fresh-frozen tissue sections.

## CONCLUSION

In summary, SpatialDecon demonstrated the most reliable and consistent performance across both simulated and experimental conditions, providing a robust framework for accurate cross-platform transcriptomic deconvolution. Under challenging settings (e.g. smaller gene panels), the methods tends to underestimate differences in cell type proportions, which may lead to false-negative results but helps to avoid the identification of false-positive cell-type differences. Overall, transcriptional deconvolution and comparison across technological platforms are feasible for medium- to large-sized samples and regions of interest but become unreliable for small cell populations comprising fewer than ca. 75 cells.

## SUPPLEMENT

### Supplementary Figures

**Figure S1:**
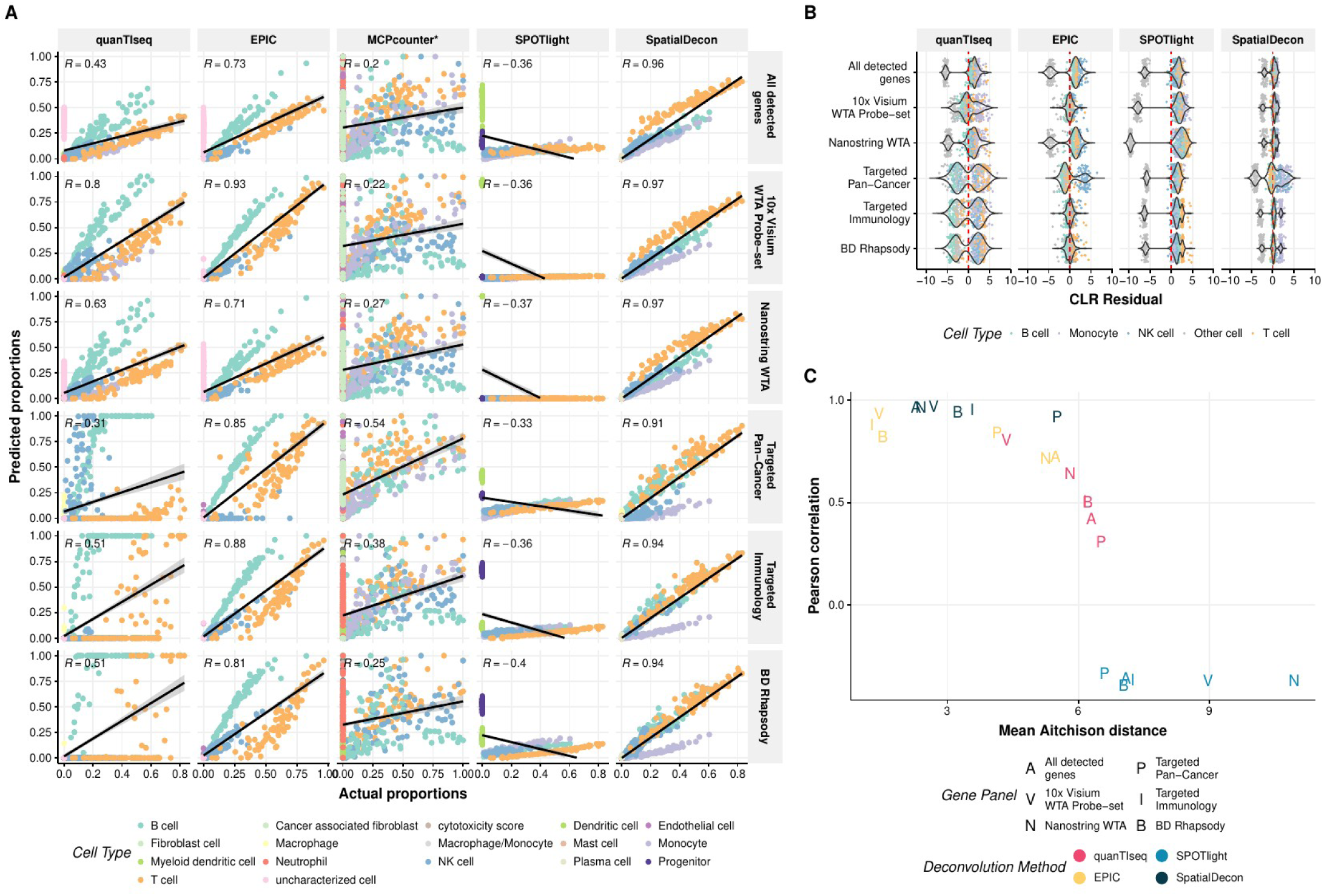
Impact of gene panel on deconvolution performance in bone marrow pseudo-bulk samples. **(A)** Correlation plots between actual and predicted proportions of cells in bone marrow pseudo-bulk samples. The colors of the dots represent the different cell types. Each plot facet displays R, which is the PCC. **(B)** Distribution of CLR residuals by deconvolution method and gene panel. The color of the dots denotes the cell type. **(C)** PCC versus mean AD for the performance of gene panels on the deconvolution tools in the PBMC pseudo-bulk samples.

**Figure S2:**
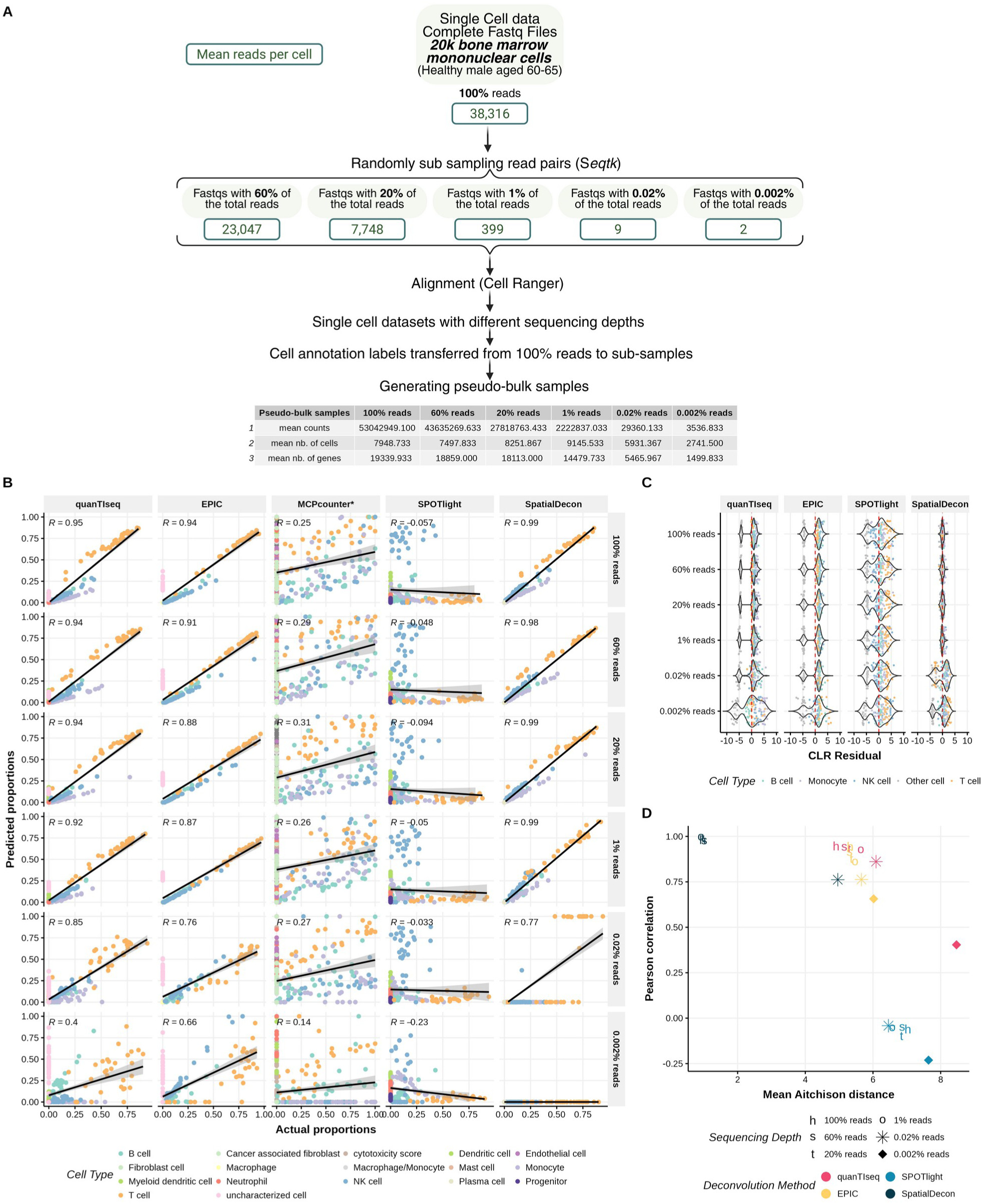
Impact of sequencing depth the deconvolution performance in bone marrow pseudo-bulk samples. **(A)** Reads were randomly sampled from the FASTQ files to obtain 100%, 60%, 20%, 1%, and 0.02% and 0.002% of the original total reads. CellRanger was run for each subsample, and cell annotations were transferred based on clustering from the 100% reads sample. These data were then processed following the method described in Fig. 1B to generate pseudo-bulk samples. **(B)** Correlations between actual and predicted cell proportions in the PBMC pseudo-bulk samples. Dots colors indicate different cell types. PCC is indicated in each plot. **(C)** Distribution of the CLR residuals for deconvolution results across different methods and sequencing depths. Dot colors represent cell types. **(D)** PCC versus mean AD comparing deconvolution performance across sequencing depths.

**Figure S3:**
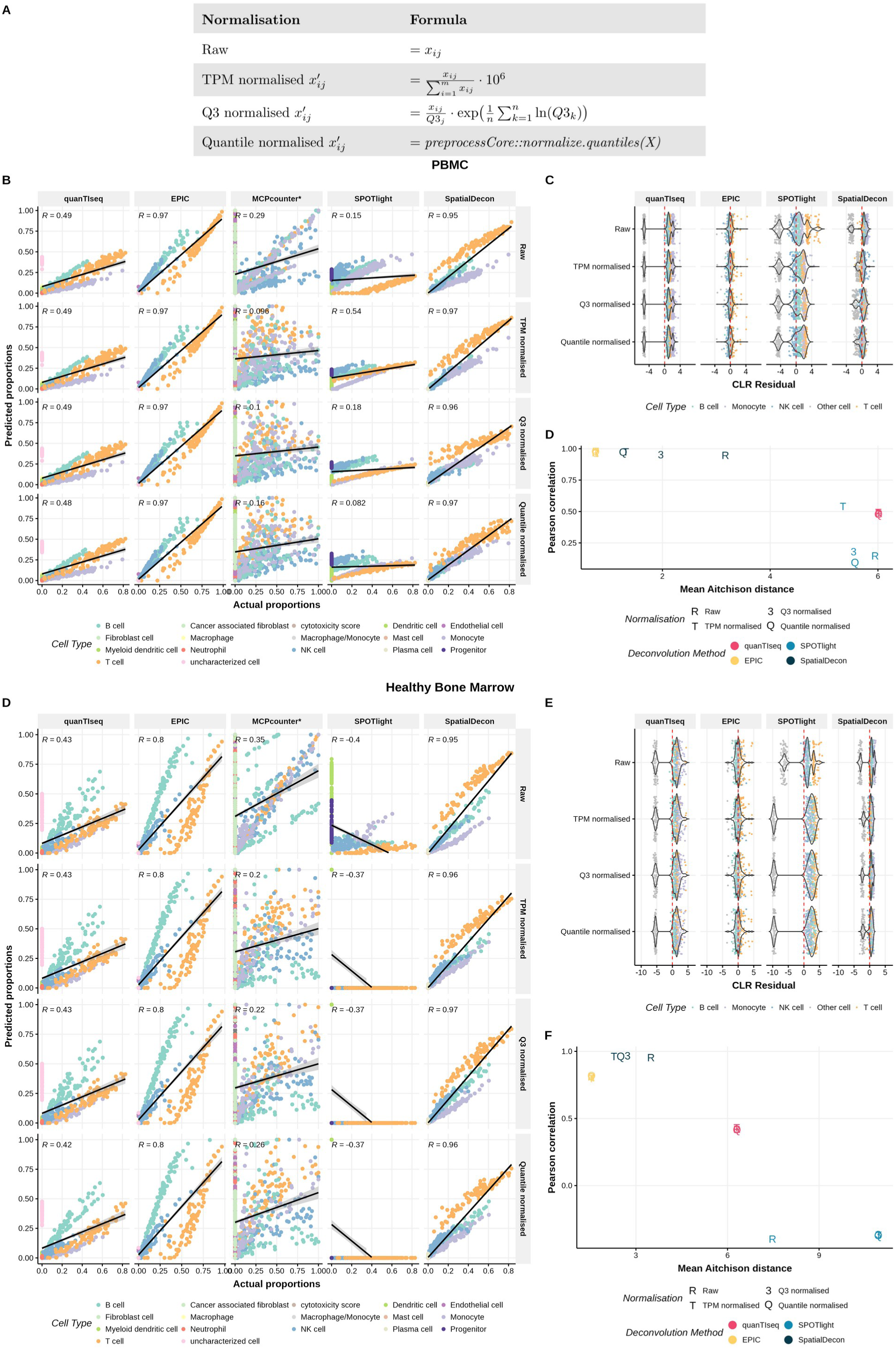
Impact of normalisation strategies on predicted cell type proportions in PBMC and healthy bone marrow deconvolution. **(A)** Formulas used to normalize pseudo-bulk samples using four different normalisation strategies: raw (no normalisation), TPM normalised, Q3 normalisation, and quantile normalisation. **(B, E)** Correlation between actual and predicted proportions of cells in the PBMC **(B)** and healthy bone marrow **(E)** pseudo-bulk samples. Each plot facet displays the PCC. Data were normalised using strategies mentioned above. **(C, F)** Distribution of CLR residuals for the four cell types in PBMC **(C)** and healthy bone marrow **(F)** pseudo-bulk samples. The color of the dots denotes the cell type. **(D, G)** PCC versus mean AD comparing different combinations of normalisation methods and deconvolution tools in PBMC **(D)** and healthy bone marrow **(G)** pseudo-bulk samples.

### Supplementary tables

**Supplementary Table 1. Summary of datasets used in this study.**

Includes dataset descriptions, GEO accession codes, and links to publicly available data.

**Supplementary Table 2. Summary of Cell-Type Classifications Across Methods**

Lists the cell types identified from single-cell data and predicted by each deconvolution tool, along with their grouping into broad cell-type categories used in the analysis.

**Supplementary Table 3. Genes included in each panel.**

Lists of HGNC gene symbols from five gene panels: all detected genes with HGNC names, 10x Visium Human Transcriptome Probe Set 1, Nanostring WTA, custom BD Rhapsody panel, and 10x Genomics Xenium Human Immuno-Oncology panel.

## MATERIAL AND METHODS

### Cross-platform datasets included in the study

Details of all the datasets used in the study are provided in Supplementary Table 1.

#### 10x Visium Data from Human Ovarian Cancer (publicly available)

10x Genomics Visium Human Ovarian Cancer datasets with different gene panels (Dataset WTA, TI, TPC as listed in Table S1) were accessed using the TENxVisium package^44^ from Bioconductor.

#### Nanostring GeoMx DSP from Human Glioma (generated in-house)

Nanostring GeoMX DSP data from glioma samples, including matching mIF image data (for 28 ROIs from 10 patients), were generated in-house. These datasets, along with their processing, are detailed in Cakmak et al. (2025)^1^ and are available under GEO accession ID (GSE271255). Two BM samples were also processed using the same protocol, the data from these samples is available upon request from the authors.

#### 10x Visium Data from Human Glioma and Brain Metastasis (generated in-house)

Four samples (2 Glioma and 2 brain metastasis) processed with Nanostring GeoMX DSP platform were also analysed using the 10x Genomics Visium protocol to enable cross-platform comparison. FFPE tissue sections (5 µm thick) were placed onto a Visium Spatial Gene Expression slide and heated at 42°C for 3 hours on a thermocycler using the Visium PCR Adaptor. The slides were then processed according to the manufacturer’s protocol for the Visium Spatial Gene Expression for FFPE workflow (CG000160, CG000239, CG000407; 10x Genomics, California, USA).

Briefly, slides underwent deparaffinisation followed by hematoxylin and eosin (H&E) staining. Stained slides were imaged using the PhenoImager HT (Akoya Biosciences, Massachusetts, USA). After imaging, coverslips were removed, and the slides were subjected to decrosslinking, probe hybridization, probe ligation, probe extension and library construction. Libraries were sequenced with paired-end dual-indexing (28 cycles Read 1, 10 cycles i7, 10 cycles i5, 90 cycles Read 2) on a NextSeq 1000 sequencer (Illumina, California, USA).

Reads were aligned to the human genome reference GRCh38-2020-A using the Space Ranger 2.0.0 pipeline (10x Genomics), and gene expression counts were quantifided. Immune-rich regions were annotated based on CD20 expression using Loupe Browser (version 6.2, 10x Genomics). The gene expression data from these samples are available upon request from the authors.

#### Deconvoluting Visium and Nanostring data

For the 10x Visium datasets, only spatial barcodes classified as ‘immune-rich’ based on CD20 expression were used for comparison with ‘immune-rich’ regions identified by the Nanostring GeoMx DSP in human glioma and brain metastasis samples. Each Visium spot and GeoMx ROI was normalised using a TPM method. To reduce platform-specific bias, only genes shared between the two platforms were retained for downstream deconvolution analyses.

#### Comparing Transcriptional deconvolution of Nanostring data to mIF proteomic cell counts

For comparison of the 28 ROIs from 10 glioma patients, the ‘immune rich regions’ from Nanostring GeoMx DSP was analyzed and matched to mIF data. The corresponding gene expression count matrix was TPM-normalised, and all genes from the Nanostring WTA panel were included for deconvolution using SpatialDecon (signature matrix= safeTME, bg=0.01).

### Generation of simulated datasets from single-cell transcriptomic data

#### Single-Cell sequencing datasets (publicly available)

We analysed several publicly available single-cell RNA-sequencing datasets. Four 10x Chromium PBMC samples (Dataset A, B, C and D; Table S1) were downloaded using the Bioconductor package TENxPBMCData^45^. Datasets A and C originate from the same patient, while Datasets B and D are from another patient. Three additional 10x Chromium bone marrow datasets (Dataset E, F, G; Table S1) from three distinct healthy donors were obtained from Bailur et al. (2020)^46^. All the above-mentioned datasets had already been pre-processed with Cell Ranger^37^ (Zheng et al., 2017) in their original studies. Dataset A and C were previously processed on Cell Ranger version 1.1.0 with hg19 transcriptome. Dataset B and D were previously processed on Cell Ranger version 2.1.0 with GRCh38 transcriptome with Single Cell 3’ v2 chemistry. Dataset E, F and G were previously processed on Cell Ranger version 1.2.0 with hg38 transcriptome.

For our sequencing depth analysis, we retrieved the raw FASTQ files of PBMC20k dataset (Dataset H; Table S1) and 20K bone marrow dataset (Dataset I; Table S1), from the 10x Genomics website.

#### Cell-type annotation

Data from each origin (PBMC or bone marrow) were compiled into SingleCellExperiment objects^47^. Preprocessing included the removal of cells identified as outliers based on mitochondrial gene content. Cells were clustered using the k-nearest neighbour graph (k = 10), using the ‘scran’ package^48^. Cell type annotation was performed using SingleR^26^, with the MonacoImmuneDataset^24,25^ from the celldex package as the reference. For consistency in downstream analysis, we grouped fine-grained cell types into broader categories, e.g. CD8+ T cells and CD4+ T cells were labeled as “T cells”. A summary of the cell types identified in single-cell datasets and their grouping into broad cell-type categories is provided in Table S2.

#### Generation of simulated bulk samples

Random numbers of each of the four cell types (B cells, T cells, monocytes, and NK cells; Monocytes were excluded from the pseudo-bulk samples used in EPIC) were sampled from their respective sub-populations within the processed single cell dataset, ensuring a minimum of 10 cells per cell type in each pseudo-bulk sample. For example, T cells were sampled exclusively from the T cell population, B cells from the B cell population, and so forth. To assess the impact of the maximum number of cells per sample, minimums cell counts of 2, 25, and 200 cells per cell type were used. Raw count expression values were aggregated to generate pseudo-bulk samples. To reduce sampling bias and improve statistical robustness, the sampling process was bootstrapped 30 times for each single-cell dataset, creating replicates and capturing intra-cellular heterogeneity in each sample. The actual numbers of cells of each cell type used in each pseudo-bulk sample was recorded during this process.

### Simulation of different gene panels

We tested five commonly used gene panels using pseudo-bulk data:

a. 10x Visium Human Transcriptome Probe Set (v1.0)^31^: Contains 19,144 genes targeted by 19,902 probes.
b. Nanostring GeoMX Human Whole Transcriptome Atlas^30^: Includes 18,816 genes.
c. 10x Visium Targeted Immunology^28^: Contains 1056 genes.
d. 10x Visium Targeted Pan Cancer^29^: Contains 1253 genes.
e. BD Rhapsody Immune Response Panel: A custom-designed primer panel was added to the predesigned human Immune Response Targeted Panel (BD Biosciences, cat# 633750) to amplify the cDNA of 434 different genes. This gene panel is published in Bexte et al, (2024)^27^.

As a positive control, a panel containing all genes detected in the dataset (that have HUGO approved nomenclature) was used (22,649 for PBMC and 33,514 for healthy bone marrow). This approach allowed us to evaluate the effect of gene panel choice across commonly used panels. Details of the genes included in these panels are provided in Supplementary Table 2.

Pseudo-bulk samples were subset based on the gene panel being tested, and deconvolution pipelines were run to predict cell types. For other parameter tests, the default panel (all genes) was used.

### Simulation of Sequencing Depth

Using the “PBMC20k” (Dataset H in Table S1) and “20k bone marrow mononuclear cells” (Dataset I in Table S1) datasets, we simulated reduced sequencing depths by downsampling the original reads to 60%, 20%, 1%, 0.02% and 0.002% using seqtk^49^. The resulting FASTQ files were aligned with Cell Ranger (version 9.0.0)^37^, using the GRCh38-2024-A reference genome. Cells in the full dataset (100% reads) were annotated following the protocol described in the ‘Single-Cell Pipeline’ section. These annotations were transferred to the downsampled datasets to ensure reliable and consistent cell type labelling across all sequencing depths, as annotations at very low read depths (e.g., median ∼2 reads/cell at 0.002%) tend to be unreliable. Finally, pseudo-bulk samples were generated by aggregating cells of the same annotated type, following the previously described workflow.

### Simulation of different cell numbers per sample

We evaluated three spot sizes to study the effects of sample size on deconvolution:

a. 6-12 cells per spot: Mimics 10x Visium spots, includes 2-3 cells of each cell types tested in the analysis.
b. 75-496 cells per spot: Represents Nanostring GeoMx DSP regions, with a minimum of 25 cells and a maximum of 124 cells per cell type.
c. 600-3996 cells per spot: Emulates larger tissue sections; containing between 200 and 999 cells per cell type.

For other analyses, we used a minimum of 10 cells and up to the maximum available cells per type in each dataset.

### Transcriptional deconvolution

#### Tools used in this study

We applied five deconvolution tools to estimate the cell composition of the samples:

a. quanTIseq^20^: Uses constrained least-squares regression with TIL10 as the default signature matrix.
b. EPIC^21^: Based on least-squares regression and includes BRef and TRef signature matrices. TRef is used as the default.
c. MCPcounter^22^: Estimates cell-type abundance using the log2 geometric mean of predefined marker gene sets. The output scores are best suited for comparing a given cell type across samples but not directly comparable between cell types within the same sample. Therefore, we scaled the predicted scores for each cell type to the [0,1] interval across samples using the following formula:

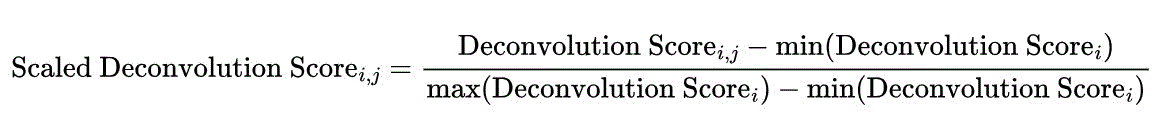

where *i* is the cell type and *j* is the pseudo-bulk sample. This allowed the preservation of the relative distribution of each cell type across samples. Actual cell counts in the pseudo-bulk samples were scaled using the same procedure to allow meaningful comparison with the predicted values. The above mentioned tools were accessed using the ImmuneDeconv R package^50^, which provides a unified interface for multiple bulk RNA-seq deconvolution methods.
d. SPOTlight^32^: Uses seeded non-negative matrix factorization (NMF) regression and requires a single-cell reference to build the signature matrix. For PBMC pseudo-bulks samples, Dataset D was used as the single-cell reference, while Dataset F was used for healthy bone marrow pseudo-bulks. For sequencing depth analysis, the 100% PBMC20k and 100% Bone Marrow20k datasets were used as references.
e. SpatialDecon^23^: Employs log-normal regression and includes the SafeTME signature matrix as the default.

Details of the cell types detected by each deconvolution tool are provided in Table S3.

#### Different normalisation approaches

We evaluated four different normalisation methods for the pseudo-bulk datasets: Raw counts: Unnormalised gene expression values.

(a) TPM (Transcripts per Million): Gene counts divided by the total count of all genes in the pseudo-bulk sample, scaled to one million.

(b) Q3 Normalisation: Counts normalised by dividing each gene’s value by the 75th percentile (Q3) value within the sample.

(c) Quantile Normalisation: Normalizes within rows and columns by ranking genes and calculating a row-wise average per gene.This was performed using the preprocessCore package^51^.

All pseudo-bulk datasets were normalised simultaneously under each method, and the deconvolution tools were run to predict cell-type composition. TPM was used as the default normalization method for all other analyses unless otherwise specified.

#### Signature matrices

We evaluated four published immune cell signature matrices:

a. *ImmunoStates*^38^: Comprise 319 genes representing 20 immune cell types, created using diverse PBMC datasets to capture broad signatures.
b. *LM22*^39,40^: Includes 547 genes for 22 cell types. This matrix is commonly used with the CIBERSORT tool.
c. *TIL10*^21^: Consists of 170 genes for 10 cell types. It is the smallest matrix tested and is used by quanTIseq.
d. *SafeTME*^23^: Contains 906 genes across 18 cell types. It is the default matrix for SpatialDecon.

In addition to these, we generated custom signature matrices using a reference dataset corresesponsing to each pseudo-bulk dataset. These matrices included all genes from the pseudo-bulk data and cover all identifiable cell types in the reference. SpatialDecon was used to test these matrices, as it supports both single-cell and user-defined signatures. Unless otherwise stated, each deconvolution tool was run with its default matrix.

### Benchmarking metrics

#### Pearson correlation coefficient

Pearson’s R values were calculated between the actual and predicted proportions. Theses were calculated while plotting using the stat_cor() function from the R package ggpubr R package^52^ and visualised on correlation plots.

#### Aitchison Distance (AD)

Aitchison Distance was used to measure the compositional differences between the actual and predicted cell-type proportions. For a given pseudo-bulk samples, AD was calculated using the following formula:

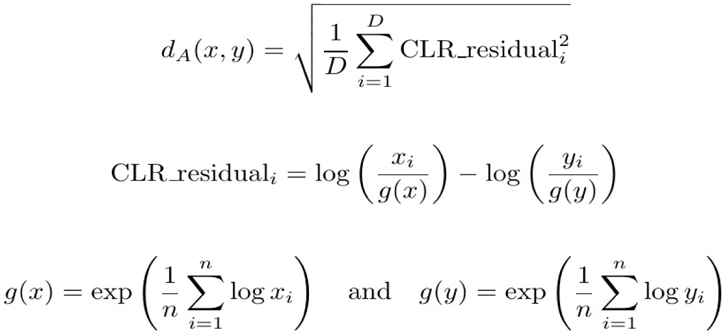

Where:

- *d _A_* is Aitchison Distance for a pseudo-bulk sample
- *x_i_* and *y_i_* are the actual and predicted proportion of a cell-type in a pseudo-bulk sample
- g(x) is the geometric mean of X (actual proportions) and g(y) is the geometric mean of Y (predicted proportions).
- D is the number of components in a pseudo-bulk sample, i.e. number of cells in a pseudo-bulk sample.

The actual compositions included only four cell types (B cells, T cells, Monocytes, NK cells; except for EPIC which excluded Monocytes), while some deconvolution tools predicted additional cell types. To address this, all the additional predicted cell types were grouped as ‘Other cells’. This ensured both actual and predicted compositions had the same number of components and preserved the Atchison geometry.

To handle discrepancies:

- If a cell type was predicted but not present in the actual data, its actual proportions was set to 0.
- If a cell type was present in the actual data but not predicted, its predicted proportion was set to 0
- A small offset (0.001) was added to all proportions to avoid zeros, since the AD operates on CLR transformedvalues, which require a positive input^35^.

Mean AD was calculated by averaging the AD across all pseudo-bulk samples of a specific experimental subfactor.

### Generating plots and figures

All plots were generated using ggplots^53^ and multi-panel figures were assembled using ggpubr^52^. Fig 1A, 2A, 3A, 4A and the graphical abstract were created using BioRender.

## DATA AVAILABILITY

All scripts used for data processing, normalization, deconvolution, benchmarking the mentioned experimental factors and generating the figures in the manuscript are available in the Github repository: https://github.com/AGImkeller/crossplatform_deconvolution.

## AUTHOR CONTRIBUTION

A.S. and K.I. conceived the project and developed computational methodology. A.S. performed computational experiments, analyzed data, and generated figures. J.H.L., J.S., P.C., and J.M. performed laboratory experiments. K.H.P and Y.R. supervised laboratory experiments. K.H.P. provided sample material. A.S. and K.I. wrote the manuscript. All authors reviewed and approved the final manuscript.

## ACKNOWLEDGEMENT

The authors would like to thank Jonas Schuck and Charles Gwellem Anchang for helpful feedback on the manuscript. K.I. was supported by funding from the Mildred Scheel Career Center Frankfurt (Deutsche Krebshilfe). K.I. and A.S. were funded by the Frankfurt Cancer Institute (LOEWE program). J.H.L. is supported by the Uniscientia foundation. P.C. was supported by the Edinger foundation. The Immunomonitoring Platform (K.H.P.) is supported by Frankfurt Cancer Institute (LOEWE program), German Consortium for Translational Cancer Research (DKTK), Uniscientia foundation (Vaduz) and Ludwig Edinger foundation (Frankfurt). Biorender was used to create figure Fig 1A, 2A, 3A and 4A.

