## Supplementary Table 2 for "Benchmarking cell-type deconvolution in cross-platform transcriptomic data"

| **Cell-Type as used in the Analysis** | **Cell annotations in single-cell data generated using the Monaco Immune Data** | **quanTIseq**  (TIL10 signature) | **EPIC**  (TRef signature) | **MCPcounter**  (Default marker list) | **SpatialDecon**  (SafeTME signature) |
| --- | --- | --- | --- | --- | --- |
| B cell | B cells | B cell | Bcells | B cell | B.Naive  B.Memory |
| T cell | CD4+ T cells  CD8+ T cells  T cells | T cell CD4+ (non-regulatory)  T cell CD8+  T cell regulatory | CD4_Tcells  CD8_Tcells | T cells  T cell CD8+ | T.CD4.Naive  T.CD4.Memory  T.CD8.Naive  T.CD8.Memory  Treg |
| NK cell | NK cells | NK cells | NKcells | NK cell | NK |
| Monocyte | Monocytes | Monocyte |  | Monocyte | Monocytes.C  Monocytes.NC.I |
| Other (Myeloid) | Dendritic cells  Basophils  Neurtrophils (not detected in single cell datasets used in this analysis) | Macrophage M1  Macrophage M2  Myeloid Dendritic cell  Neutrophil | Macrophages | Macrophage/Monocyte  Myeloid dendritic cell Neutrophils | mDCs  pDCs  Neutrophils.LD  Macrophages |
| Other (Stromal) |  |  | Endothelial  CAFs | Endothelial cell  Cancer Associated Fibroblasts | Gentles.endothelial Gentles.fibroblasts |
| Others (Misc) | Progenitors | uncharacterized cell | otherCells | Cytotoxicity score | Mast.cells  Plasmablasts |

**Supplementary Table 2:** Summary of cell types identified in the single-cell data and by deconvolution tools, along with their grouping into the broad cell-type categories used in the analysis
